# Unsupervised modeling of mutational landscapes of adeno-associated viruses viability

**DOI:** 10.1101/2023.10.26.564138

**Authors:** Matteo De Leonardis, Jorge Fernandez-de-Cossio-Diaz, Guido Uguzzoni, Andrea Pagnani

## Abstract

Adeno-associated viruses 2 (AAV2) are minute viruses renowned for their capacity to infect human cells and akin organisms. They have recently emerged as prominent candidates in the field of gene therapy, primarily attributed to their inherent non-pathogenic nature in humans and the safety associated with their manipulation. The efficacy of AAV2 as gene therapy vectors hinges on their ability to infiltrate host cells and subsequently replicate within them, a phenomenon reliant on their competence to construct a capsid capable of breaching the nucleus of the target cell. To enhance their infection potential, researchers have extensively scrutinized various combinatorial libraries by introducing mutations into the capsid, aiming to boost their effectiveness. The emergence of high-throughput experimental techniques, like Deep Mutational Scanning (DMS), has made it feasible to experimentally assess the fitness of these libraries for their intended purpose. Notably, machine learning is starting to demonstrate its potential in addressing predictions within the mutational landscape from sequence data. In this context, we introduce a biophysically-inspired model designed to predict the viability of genetic variants in DMS experiments. This model is tailored to a specific segment of the CAP region within AAV2’s capsid protein. To evaluate its effectiveness, we conduct model training with diverse datasets, each tailored to explore different aspects of the mutational landscape influenced by the selection process. Our assessment of the biophysical model centers on two primary objectives: (i) providing quantitative forecasts for the log-selectivity of variants and (ii) deploying it as a binary classifier to categorize sequences into viable and non-viable classes.

## 1. INTRODUCTION

Recently, there has been a burgeoning interest in combining high-throughput sequencing with machine learning techniques to extend predictions beyond the boundaries of experimentally observed sequences Wu et al. (2019). While previous studies predominantly concentrated on a single protein property directly associated with the selection criteria, such as binding, stability, or catalytic activity Araya et al. (2012), a few investigations have showcased the feasibility of inferring multiple physical properties, including those that are not directly measurable Kinney & McCandlish (2019). Notable examples of success include predicting thermal stability based on binding affinity measurements Otwinowski & Plotkin (2014); Otwinowski et al. (2018) and deducing specificity profiles of transcription factors from the selective enrichment of DNA sequences Rastogi et al. (2018); Rube et al. (2022). The key to these achievements lies in the application of biophysics-inspired models, which incorporate thermodynamic principles to yield interpretable predictions.

The combination of high-throughput *in-vitro* selection methods with high-throughput sequencing has been key to identifying protein variants of a given wild-type sequence with target functional activity Fowler & Fields (2014); Boyer et al. (2016); Schulz et al. (2021). However, it’s crucial to recognize that all experimental approaches are constrained by the maximum library size, typically falling within the range of 10^8^ (for yeast display), 10^10^ (for phage display), to 10^15^ (for ribosome display). Although these numbers may seem substantial, they represent only a minuscule fraction of the vast sequence space (for example, there are 20^28^ ∼ 2*×* 10^36^ possible protein sequences of length 28). Lately, there has been a significant surge of interest in directed evolution experiments (both in vitro and in vivo) aimed at engineering the capsid of adeno-associated virus (AAV) Wu et al. (2000); Dalkara et al. (2013); Tse et al. (2017); Ogden et al. (2019). AAV2 are small non-enveloped viruses of a typical dimension of about 20 nm. They are provided with a spherical capsid composed of 60 subunits arranged with icosahedral symmetry. While they can infect humans and other primate species, there are no currently known diseases that they can cause. This, besides other technical motivations, makes them ideal candidates as viral vectors in gene therapy as shown in several clinical tests.

The use of AAV2s in gene therapy is conditioned by their potential to infect cells which in turn depends on their capacity to assemble a viable capsid, meaning that they assemble an integral capsid that packages the genome. From the mechanical point of view, much interest has been attracted by the idea of understanding the assembly process of the viral capsid Mendoza & Reguera (2020), bringing considerable insights about the patterns that lead the capsid formation starting from its basic building blocks (individual capsid proteins, oligomers or capsomers). From a microscopic point of view, capsid formation can be described as a nucleation process driven by three free-energy contributions: (i) the energy gain in growing a partial shell given by the binding energy between capsid proteins, (ii) a domain wall energy penalty due to missing contacts at the border of the shell, and (iii) the elastic energy due to the curvature of the shell shape. Thanks to experimental observations and simulation analysis, it is now possible to highlight the leading factors that make the assembly possible in different mechanical conditions and also the microscopic patterns through which the capsid can assume the correct shape. A bottleneck in achieving the desired phenotypic traits of the AAV genotype is associated with capsid production, where the majority of sequence variants fail to either assemble or package their genome. This phenomenon is commonly referred to as *viability*.

Here, we address the problem of forming a viable capsid from a Deep Mutational Scanning (DMS) experiment perspective. In particular, we study how mutations of a particular section of the CAP region affect capsid formation from a macroscopic point of view. The idea of this work is to devise a new machine-learning strategy that improves currently employed computational techniques to analyze data from DMS experiments. Instead of training a deep neural network to solve a regression problem where the sequences are the input of the model and the output is the selectivity of the sequence, we aim to develop a biophysical model of a general DMS experiment, with physically interpretable parameters that remain comparable between different experimental realizations. On top of this advantage, we want to show how a biophysical model can lead to a more robust inference: less biased by experimental noise and more accurate in predicting the selectivity of the variants especially when we are interested in comparing it between different variants.

To this aim, we analyzed data collected in a massive study on capsid diversity (Bryant et al. (2021)). In this study a DMS experiment has been performed testing a large library of viruses with diverse capsid composition, these data have been then used to train different machine learning models to infer the mutational landscape for capsid viability and use it to generate viable capsids as different as possible from the WT one. The statistical model employed in our approach is an energy-based model, expressed through a combination of convolutional layers (accounting for variations in sequence lengths within the training sets) and a dense layer, responsible for mapping sequences to energy levels. Notably, the performance of this biophysics-inspired model proves intriguing even in a more straightforward classification scenario, where it is applied to the binary task of distinguishing viable from non-viable sequences.

## 2. EXPERIMENT AND DATA

The data used in our analysis are derived from a recent comprehensive study Bryant et al. (2021). In this work, the variability of protein capsids in AAV2 and their ability to remain viable for DNA packaging was investigated by developing a deep learning method to estimate the mutational landscape for capsid viability and generate new diverse viable capsids that differ significantly from those found in nature. The analysis was focused on a specific region comprising 28 amino acids near the 3-fold symmetry axis of the icosahedral AAV2 VP1 protein. This region is known to play a crucial role in viral production. The phenotypic trait selected for in the experiment is the virus viability and it was quantitatively assessed through high-throughput screening experiments to collect data on a wide range of variants. To do so, a series of libraries of designed capsid proteins were synthesized and inserted into the virus genome’s cap region. Plasmids encoding various capsid structures were cloned to create an extensive plasmid library. These plasmids were first sequenced and then transfected into cells. After viral production, the DNA was extracted and sequenced once again, resulting in two sets of reads for each tested variant. These reads provided information about the abundance of each sequence before and after transfection.

The training set for our algorithm comprises three data sets that differ considerably one from another in both size (number of unique variants) and diversity (distance from the wild-type sequence). To assess their diversity, we consider the Levenshtein distances of the sequences from the wild-type one (from now on we will always imply this particular metric whenever we talk about distance). In *experiment-1* single-site mutations and insertions concerning the wild-type sequence were tested (so all variants are at a distance 1 from the wild-type). In *experiment-2* sequences sampled at random around the wild-type up to a maximum distance were tested (see Figure 1 left panel). In *experiment-3* sequences generated using the machine learning methods employed in Bryant et al. (2021) were tested.

**Figure 1:**
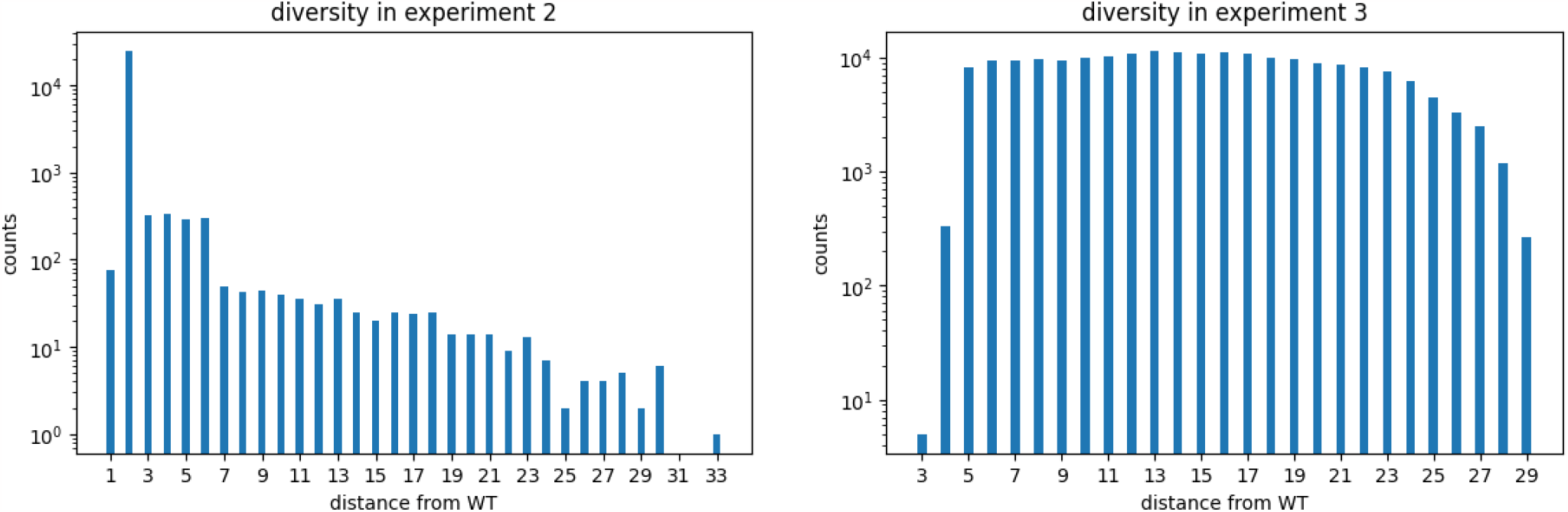
Distances of the tested sequences (from the WT sequence) in each of the two experiments performed in Bryant et al. (2021).

**Table 1:** Relevant information about size, quality, and diversity of the dataset tested in each experiment.

|  | experiment-1 | experiment-2 | experiment-3 |
| --- | --- | --- | --- |
| unique variants | 1085 | 26961 | 203635 |
| depth round 0 | 2423857 | 14353318 | 49376644 |
| depth round 1 | 3491860 | 23411926 | 194466876 |
| coverage round 0 | 2234 | 532 | 242 |
| coverage round 1 | 3218 | 868 | 955 |
| distance from WT round 0 ($\mu \pm \sigma$) | $0.9 \pm 0.2$ | $3 \pm 2$ | $13 \pm 5$ |
| distance from WT round 1 ($\mu \pm \sigma$) | $0.9 \pm 0.3$ | $3 \pm 2$ | $10 \pm 4$ |

Int Fig. 1 we display the diversity of each training set. The height of the bars refers to the number of different *unique* variants tested in each experiment. Tab. 1, instead, summarizes the main features of the three datasets.

## 3. METHODS

In this section, we describe the details of our biophysical-inspired statistical model, and describe the different training and test datasets used. The model defines a statistical *energy* that we consider as a proxy to the sequence viability. Strictly speaking, viability is a discrete phenotypic trait that in the series of experiments conducted that constitute our training set Bryant et al. (2021) is assessed in terms of sequencing counts. The boolean nature viable/non-viable is eventually induced with a statistical analysis *a-posteriori*. To compare the performance of the our model in solving the simpler boolean classification task, we also introduce a standard classifier with the same architecture as the energy-based model.

### 3.1. Model

The machine-learning method we propose utilizes a statistical model that aims to capture the entire experimental pipeline conducted to assess the ability of variants to package the genome within the capsid. The structure of this model is very general and permits the analysis of any DMS (Deep Mutational Scanning) experiment. As previously discussed in Fernandez-de-Cossio-Diaz et al. (2020), the model comprises three main steps: selection, amplification, and sequencing.

During the selection phase, mutated viruses are introduced into tissue cells, where their survival and replication depend on their ability to form functional capsids. Typically, only a small fraction of the initial population of viruses is selected during this phase. Following selection, viral DNA is extracted from a sample of cells and undergoes amplification. Finally, the amplified DNA is sequenced to gather information about the genetic composition of the selected viruses.

#### Selection

We examine a population of AAV2s (Adeno-Associated Viruses), each carrying a mutated sequence of the gene of interest. Let *N*_*s*_ represent the count of viruses containing sequence *s*in the initial library. From a biophysical perspective, we assume that the functional characteristics of the viral variant are determined by the thermodynamic states adopted by the protein products resulting from the mutated gene. The expressed protein copies of variant *s* can either bind together, leading to the assembly of a functional capsid, or their structural properties may hinder capsid formation. The energies of these two states will be denoted by 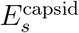 and 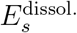, respectively. Since survival depends on the ability to form the capsid, the probability of selection for viruses carrying variant *s* (or *selectivity*) can be computed according to the Boltzmann law

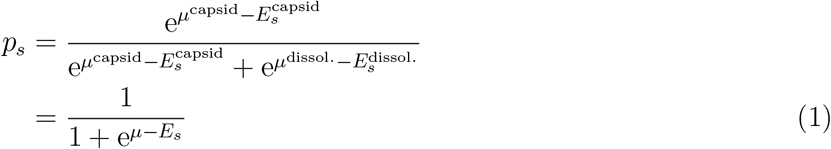

where *µ*^capsid^, *µ*^dissol.^ are chemical potentials associated with each state, and we defined *µ*:= *µ*_dissol._ − *µ*_capsid_ and 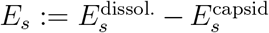. From now on, we define the pair *E*_*s*_, *µ*, energy specificity, and chemical potential respectively. With these definitions, *p*_*s*_ assumes the well-known Fermi-Dirac distribution form for a two-level system. In the large *N*_*s*_ limit, one can expect small fluctuations in terms of the number of selected sequences. Therefore, out of the *N*_*s*_ viruses carrying sequence *s*in the initial library, we assume *N*_*s*_ *p*_*s*_ are selected by their ability to form the capsid. This approximation is typically referred to as *deterministic binding approximation* Fernandez-de-Cossio-Diaz et al. (2020).

#### Energy specificity

The capsid-formation energy *E*_*s*_, depends on the ability of protein products to bind together and assemble the capsid. Protein binding depends on sequence*s*, and in full generality this dependence is complex. In analogy with the reference study, we use one of the neural network models tested in Bryant et al. (2021) to parametrize the energy mapping specificity. The network is composed of two convolutional layers and then three fully connected ones (see Sec. A.3 for the detailed structure).

We denote the set of parameters of this network (weights and biases) by Θ. Having specified the network architecture, selectivities then become functions of Θ and the chemical potential,

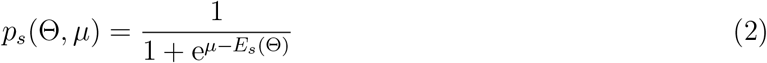

Although other choices of energy function could be used in principle, the convolutional structure chosen here allows us to handle sequences of different lengths as explained in detail in Sec. A.2.

#### Amplification

Empirically, it turns out that only a small fraction of viruses survive the selection phase. To replenish the original total population size, after each selection round an amplification phase is performed. We assume for simplicity that this process is uniform across all sequences. Then, denoting by *N*_*s*_ ′ the abundance of sequence *s*after amplification,

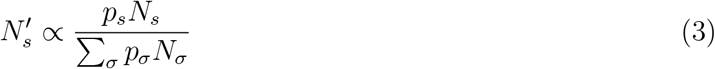

#### Sequencin

In the experimental pipeline of Bryant et al. (2021), sequencing is performed at two different steps of the experiment: (i) when the initial combinatorial library is inserted into the plasmids (but prior to transfection into the target cells), and (ii) after the viral extraction from the cell. It is clear that the target phenotype observed is the *viral viability*, i.e. the ability of the variant to remain active and capable of infecting host cells and producing progeny viruses.

If we assume that each variant is sequenced with a probability proportional to its abundance, the probability of observing a count *R*_*s*_ for variant *s* conditional to the (unobservable) number of variants *N*_*s*_, follows the multinomial distribution:

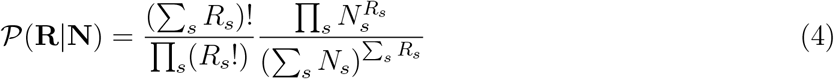

where **R**,**N**, and **p** are the vectors of read counts *R*_*s*_, variant abundances *N*_*s*_, and selectivities *p*_*s*_, respectively. Similarly, for the sample taken after selection and amplification,

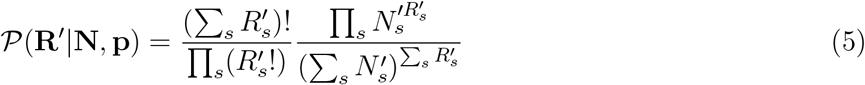

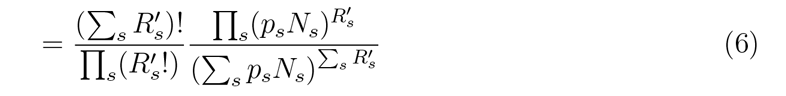

Note that *𝒫*(**R**′|**N**,**p**) depends on the selectivities *p*_*s*_, which in turn depend on the parameters Θ, *µ*, see (2).

### 3.2. Maximum likelihood inference

The model defined so far has several parameters that must be inferred from data: the parameters of the neural networks specifying the mapping from sequence to energy *E*_*s*_, that we hereby denote by Θ, the chemical potentials *µ*_*s*_, and the variant abundances in the initial library *N*_*s*_. As a function of these parameters, the model log-likelihood can be written:

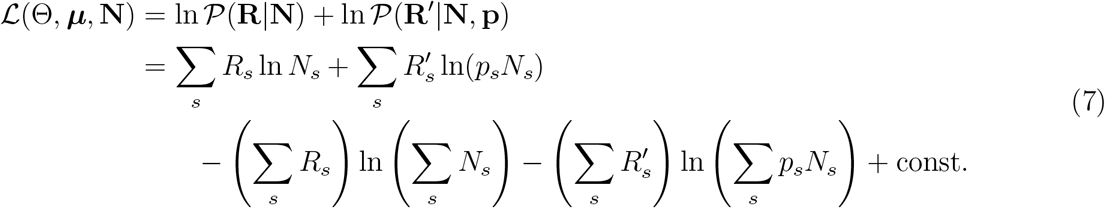

Note that the dependence on the parameters Θ, *µ* arises from the dependence of the selectivities, (2). Following Fernandez-de Cossio-Diaz et al. (2023), we then learn Θ,***µ***,**N** from the sequencing data by numerical maximization of the likelihood,

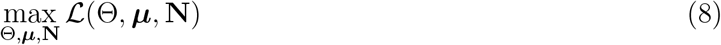

See A.4 for details on the optimization procedure.

Once the optimal parameters have been obtained, the learning can be validated by assessing how the quantities observed experimentally correlate with the ones inferred. To be more precise, we look at the correlation between *p*_*s*_ and the ratio *θ*_*s*_ = *R*_*s*_′*/R*_*s*_, usually called *empirical selectivity* Rubin et al. (2017), on a subset of sequences that are not used during training of the model. Barring sampling noise, *θ*_*s*_ should be proportional to the selectivities *p*_*s*_ inferred by the model. Unfortunately in this experiment, only one round of selection has been performed; therefore it is not possible to use information from multiple rounds to filter out experimental noise as described in Rubin et al. (2017) and already exploited in Fernandez-de-Cossio-Diaz et al. (2020).

### 3.3. Binarizing the output of the energy-based model

The most powerful feature of this machine learning method is that it is capable of inferring quantitatively the selectivity of every single variant. Anyway, there are cases in which the useful information is less specific and one can be only interested in knowing if a variant is able or not to perform its task. In these cases, one can use the model to perform a binary classification setting a threshold value for *p*_*s*_ and classifying accordingly the variants.

Let us consider the situation in which it is possible to determine experimentally a threshold value for the empirical selectivities *θ*_*t*_ and we have used the machine learning method to learn the log-selectivities *p*_*s*_. Now the goal is to find a threshold value for *p*_*s*_ so that we can classify the variants according to the learned model. What we did is the following and it consists of one possible way to find such a threshold. First, we use the inferred values of *p*_*s*_ to sort the variants assigning a permutation *π*_*s*_ ∈ {1, …, *S*} such that *π*_*s*_ ≤ *π*_*σ*_ if and only if *p*_*s*_ ≥ *p*_*σ*_. Then for each *s* ∈ *S*, we compute the fraction of empirical positive and negative sequences in the set {s′|*π*_*s*_′ ≤ *s*} according to their empirical *θ* value. Now for each *s*, we compute what is commonly known as *g-means score*,

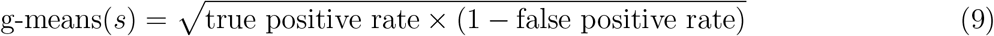

and the *s*^*^ that maximizes this score tells us the last variant*s* ^*^ that has to be considered as a positive sample, and we can choose 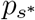 as a threshold value for the binary classification.

Consider that when we estimate the accuracy of our biophysical model, it has to be intended as a “best case” accuracy since the optimal threshold is obtained by maximizing the*g-means score* computed from the values of true and false positive rates on the test sets. For this reason, the optimal threshold is biased by a piece of information coming from the test set which, in principle, should be invisible to the classification strategy.

### 3.4. Data Binarization

We compared our biophysical model based on likelihood maximization with a more standard machine learning approach where the task here is not to quantitatively predict the log-selectivities of the selection process, but rather to perform a *simpler* binary classification of variants into the two disjoint classes of viable/non-viable. First, an empirical proxy for the “fitness” of the variants is computed, this quantity is usually referred to as *empirical (log)selectivity* and it is defined as the following

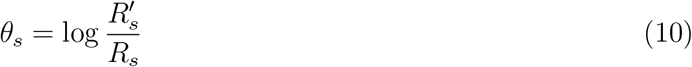

Then these quantities can be used to assign a label to each sequence; in Bryant et al. (2021) they have assigned a binary label to each variant setting a threshold for this quantity.

The procedure we used to assign this threshold value is similar to the one used in the reference study. By direct inspection of the empirical log-selectivity distribution Fig. 2, one can see that they follow a bimodal distribution. We consider sequences near the rightmost peak as viable and the remaining as non-viable. We fitted a bimodal Gaussian distribution from those histograms and we have chosen as threshold the value of log-selectivity where we find a valley in between the two peaks to have a stable classification.

**Figure 2:**
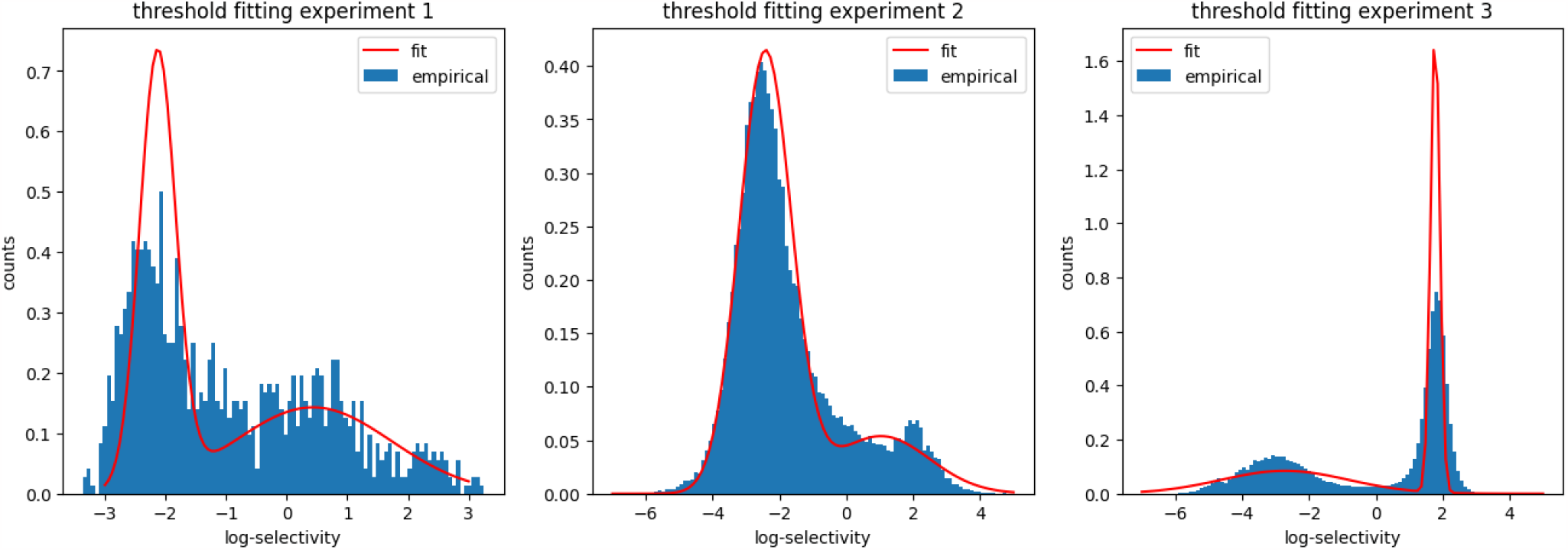
Fit performed on the histogram of log-selectivities of each experiment to determine the threshold value for binary classification of the sequences as viable or non-viable. The empirical distribution has been fitted with a bimodal Gaussian distribution and the valley in between the two peaks has been chosen as threshold value.

In contrast with Bryant et al. (2021) we decided to fit a threshold for each experiment since in principle this quantity might change for each experimental realization, and as one can see from Fig. 2 this is the case.

### 3.5. Binary classifier

To assess the classification performance of the energy-based model, we trained a binary classifier with the same architecture as the energy-based model (see Section A.2) for the details). By defining *q*_*s*,viable_ and *q*_*s*,non-viable_ respectively as the two outputs of the binary classifier associated with the two states, the loss function can be written as:

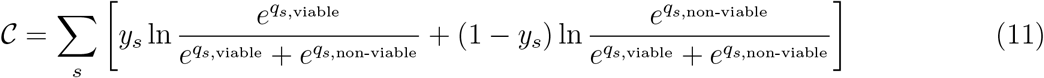

where *y*_*s*_ is 1 if the sequence is labeled as viable and 0 otherwise.

## 4. RESULTS

To test the model and its robustness we compared the two approaches, first, when we train them on single mutations/insertions sequences (those which have been tested in *experiment-1*) and use the model to infer the selectivity of sequences tested in *experiment-2*. And then we trained the models on sequences tested in *experiments 1 and 2* and used them to predict the selectivity of sequences tested in *experiment-3*.

In the first case the neural network trained on single mutations and insertions reaches an accuracy of ≈ 80% tested in *experiment-2*; instead, our biophysical model reaches a (maximum) accuracy of ≈ 84% on those sequences. Despite the similar results in terms of accuracy for binary classification, our biophysical model can provide accurate information about the relative “selectivity” between different sequences, in other terms is able to reliably infer the energy levels of the sequences. In the second case where we want to predict the selectivity of sequences in *experiment-3* we have sequences that are in principle highly diverse from the ones used in the training set. For this reason, the performance drops considerably: ≈ 67% accuracy for the neural network model and (maximum) ≈ 85% for the biophysical model. Despite the drop in the accuracy, the biophysical model is still able to maintain a high correlation between empirical log-selectivities and inferred log-probabilities of the viable state which means that the method is indeed robust when we are interested in learning the relative “fitness” (the energy levels) of the sequences, as shown in Fig. 3a and Fig. 3b (Pearson coefficient 0.75 in the first case and 0.77 in the second one).

**Figure 3:**
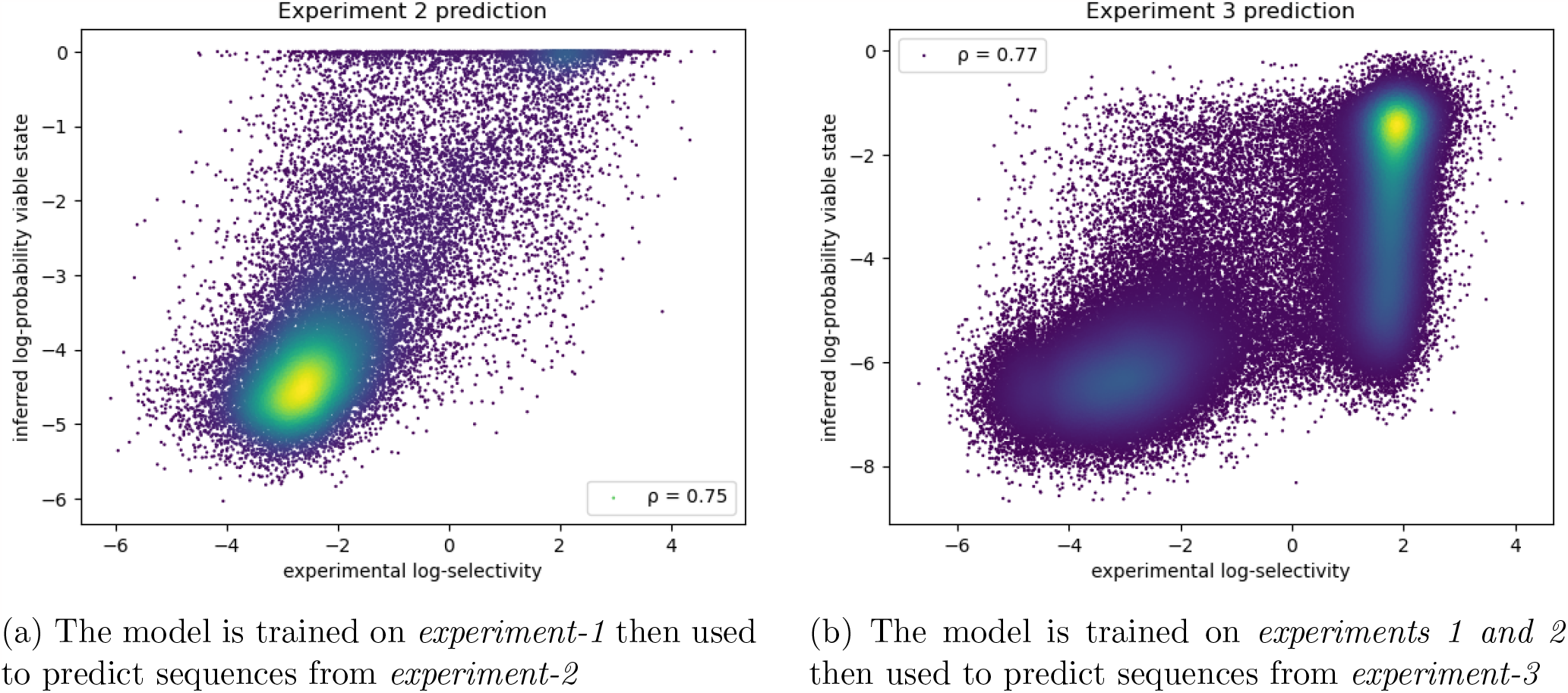
Scatter plot of empirical log-selectivities vs inferred log-probabilities of the viable state

In addition to this value of “best case” accuracy that we can provide, as described in Sec. 3.3, one can also vary the threshold value on the inferred *p*_*s*_ and realize a ROC curve to see how much the classification is robust with respect to the choice of the threshold value. A ROC curve has been realized also for the binary classifier; it has been achieved by varying the criterion used by the Neural Network to classify the sequences: instead of classifying the sequence as viable if *q*_*s*,viable_ ≥ *q*_*s*,non-viable_ we introduce a threshold*τ*and sequences are classified as viable if *q*_*s*,viable_ − *q*_*s*,non-viable_ ≥ *τ*. These two sets of ROC curves for both models are shown in fig. 4a and fig. 4b respectively for the two train/test combinations.

**Figure 4:**
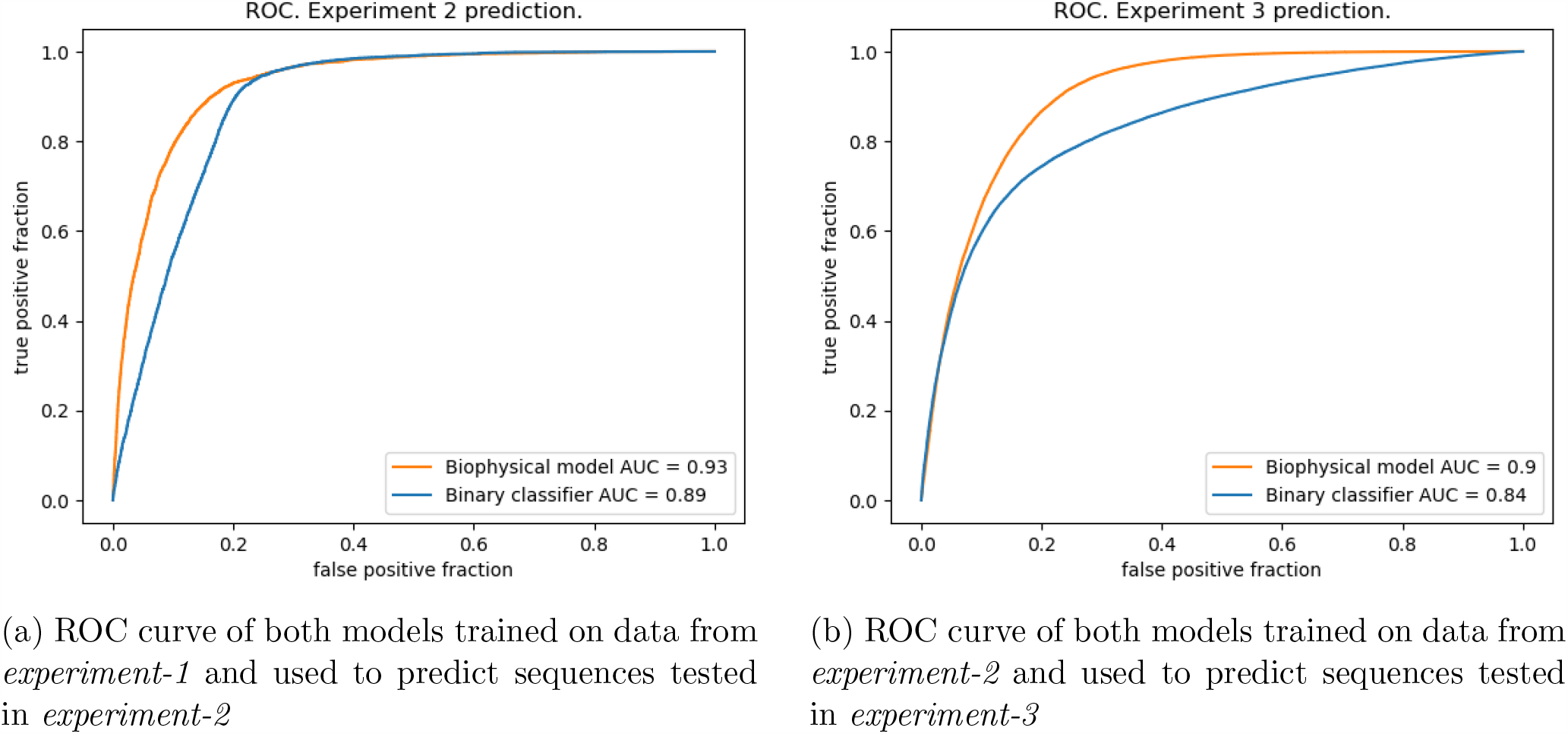
ROC curves of both models obtained by varying the classification threshold.

As one can see from Fig. 4a the area under the ROC is very high (≈ 0.93) when the model is trained on single mutations/insertions and used to test sequences from *experiment-2*. Fig. 4b shows that when the model is trained on *experiments 1 and 2* and used to predict sequences from *experiment-3* the area under the ROC slightly decreases at≈0.90. Perhaps surprisingly, we observe from Fig. 4b (orange (c) Binary Classifier trained on *experiment-1* and lines) that the performance of the classifier is marginally worse in both experiments. The same trend, perhaps more clearly, is shown in Fig. 5 where we display the confusion matrices for the two models in each train/test combination. When training both models on *experiment-1* and testing the classification task on *experiment-2*(Fig. 5 first row), we obtain a relatively comparable performance: a true positive rate of 0.90 for the biophysical model, while 0.94 for the binary classifier; a true negative rate of 0.84 for the biophysical model and of 0.77 for the binary classifier. Things change when we train on *experiment-1 and 2* and we test on *experiment-3* (Fig. 5 second row). Here, with a true positive rate of 0.44 for the binary classifier, and 0.89 for the biophysical model, we double the figure while keeping a true negative rate roughly comparable (around 0.95 for the binary classifier vs. 0.78 for the biophysical model).

**Figure 5:**
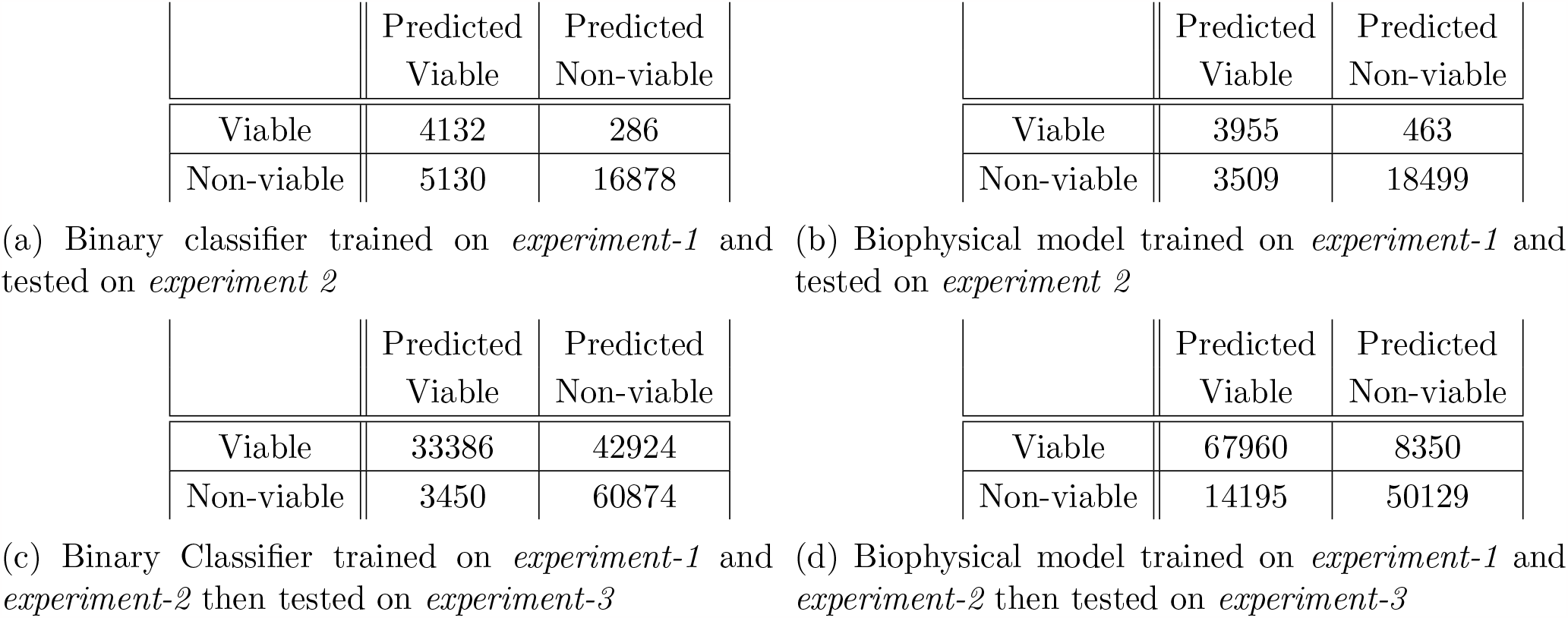
Confusion matrices of the two models for each train/test combination.

Evaluating the generalization capabilities of our model in relation to the variant distance from the WT sequence is a compelling endeavor. This test holds particular interest due to the substantial heterogeneity in composition among the various training sets, as illustrated in Figure 1. To elucidate further, *experiment-1* provides a relatively narrow view of the mutational landscape around the WT, encompassing only single inserts and mutations. Conversely, *experiment-2* and *experiment-3* explore a broader range of the mutational landscape, generating sequences at more substantial distances from the WT. Notably, *experiment-2* exhibits a higher concentration of sequences around the WT, with an exponential distribution tail extending up to a Hamming distance of 33. In contrast, the composition in *experiment-3* is more uniformly distributed across a similar range as explored by *experiment-2*. To evaluate the predictive capabilities of our models with respect to the distance from the WT sequence, we dissect the performance of the various models based on the distance of the test set sequences from the WT.

Again, we report the results for the model trained on *experiment 1* and tested on *experiment-2* (Fig. 6a and 6c) and for the model trained on *experiment-1,2* and tested on *experiment 3* (Fig. 6b and 6d). As a score, we use: the accuracy for the binary classifier, and the Pearson correlation coefficient between the model log-likelihood and the sequence empirical log-selectivity for the biophysical model.

**Figure 6:**
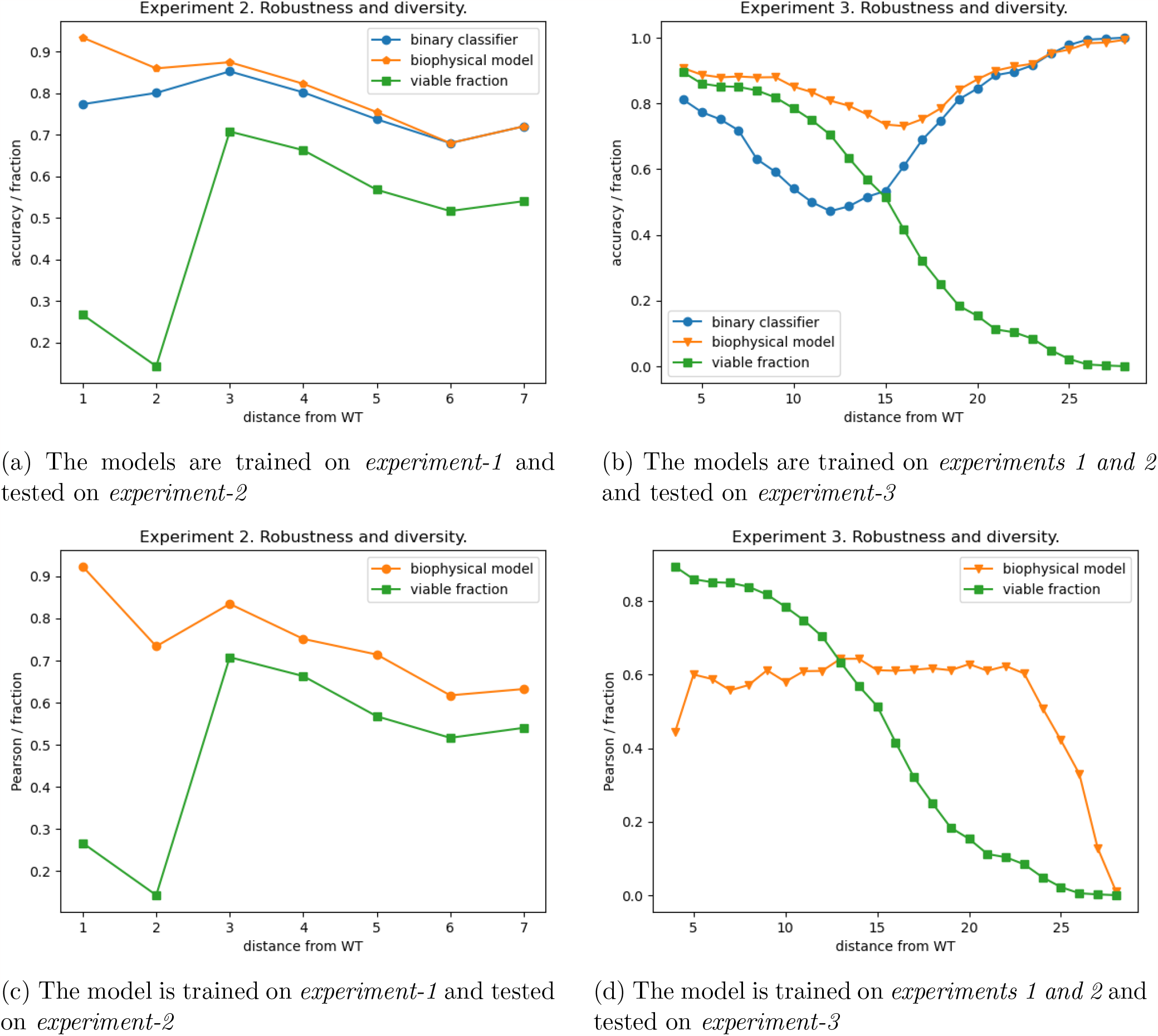
Robustness analysis with distance. The data set has been divided into slices according to their distance from wild-type and the models have been tested on those slices. The score for the two models is reported as a function of the distance of the test data.

From the green lines with square markers in Fig. 6 we immediately note how the viable fractions of the two test sets are remarkably different. Whereas *experiment-2*(Figs. 6a and 6c) shows a low fraction of viable variants at a small distance from the WT (Hamming distance lower equal than 2), this fraction increases around 0.6 at higher distances. The composition is radically different for *experiment 3* (Fig. 6b and 6d) where the viable fraction monotonously decreases as a function of the Hamming distance from the WT, from an initial value of around 0.9 to almost 0 for the most distant variants. When we compare the performance of the binary classification on the two datasets, for both models we observe that the biophysical model systematically outperforms the binary classifier although the margin is minimal when we consider as a test set *experiment-2* Figure 6a, more pronounced for *experiment-3* Figure 6b. In this second case, there is a tiny drop in performance at the intermediate distance from the WT sequence where viable and non-viable sequences are in almost equal shares, and arguably the classification task is more difficult.

Interestingly, when we assess the generalization performance of the biophysical model to predict the sequence log-selectivity when testing on *experiment-2* (Figure 6c) we observe a mildly monotonously decreasing curve from very high correlation values (around 0.9 for the closest variants) down to a correlation of around 0.7 for the most distant variants. A different behavior is observed when testing on *experiment-3*(Figure 6d)), where, besides a decrease in correlation for the extreme values of the distance from the WT, a stable correlation of around 0.6 is observed.

In spite of the overall quantitatively and qualitatively good generalization properties of our models shown by the above-mentioned results, a prominent challenge that our biophysical model still grapples with is the task of learning discernible patterns from the viable sequences examined in *experiment-3*. As shown from Figure 3b, it’s apparent that the model tends to produce false negatives. A plausible explanation for this observation becomes evident when we compare the patterns of true positives and false negatives with those of genuinely non-viable sequences. Figures 7b and 7c exhibit a strikingly similar composition, and we aspire for our model to correctly categorize all such sequences as viable, given their apparent similarities. However, when we contrast these patterns with those in Figure 7a, which represents the sequences used for training, we can discern that the model has been exposed to significantly different patterns than those we require it to recognize, particularly in the rightmost portion of the sequences. On the other hand, the model generally correctly classifies non-viable sequences because they manifest entirely distinct patterns, particularly in the leftmost section of the sequence, as illustrated in Figure 7d.

**Figure 7:**
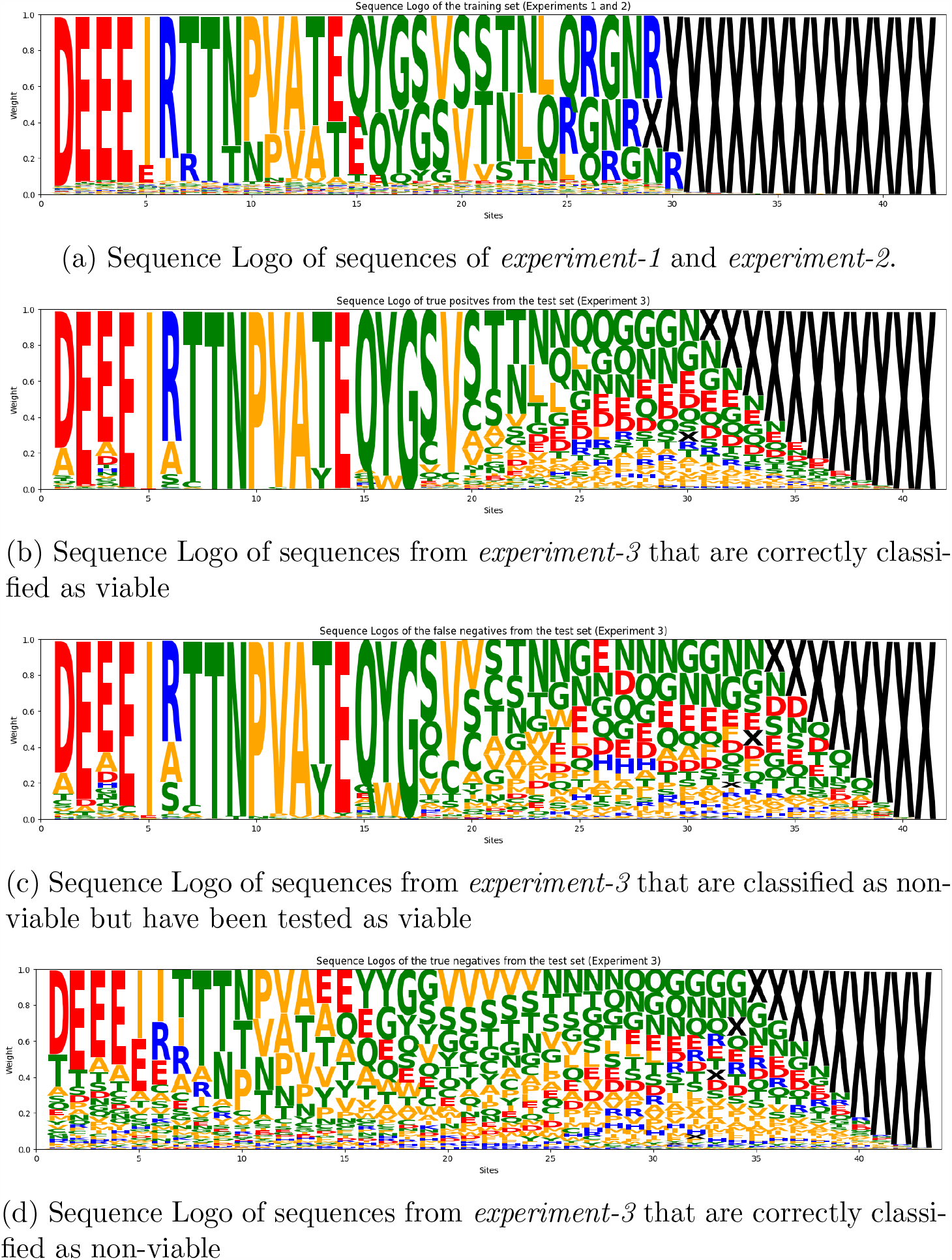
Sequence logos of some subsets of the tested sequences. The numbers on the *x* axis indicate the position along the sequence, starting from site 561 which corresponds to position 1. Those sites that are composed only of gaps within the specific subset have been cut out from the logo (for the details about the sequence encoding see Sec. A.2). The height of each letter is proportional to the frequency of the corresponding amino acid in the specific sub-set of unique sequences. The color, instead, is related to the physico-chemical characteristics of the amino acids. In some of the panels, at the rightmost sites, the gaps are so much more frequent that it is difficult to see the other small letters appearing.

## 5 CONCLUSIONS

In our research, we present a novel biophysically-inspired model designed to forecast the viability of genetic variants within the framework of Deep Mutational Scanning (DMS) experiments, as detailed in Bryant et al. (2021). Our model was meticulously fine-tuned to concentrate on a specific segment within the CAP region of the capsid protein found in Adeno-associated virus 2 (AAV2), a choice made to hone its predictive power. To gauge the effectiveness of this model, we trained our model with diverse datasets, each strategically engineered to delve into distinct sectors of the mutational landscape influenced by the selection process. Our exploration of the biophysical model’s generalization capabilities centered on two objectives: (i) furnishing quantitative forecasts for the log-selectivity of variants and (ii) deploying it as a binary classifier, categorizing sequences into viable and non-viable classes. To further enhance our model’s ability to distinguish between viable and non-viable outcomes, we developed a parallel binary classifier. This classifier adopts the same architectural framework as the biophysical model but undergoes supervised training, reinforcing its ability to classify sequences with precision.

A noteworthy feature of the DMS library presented in Bryant et al. (2021), is its incorporation of not only mutations but also a substantial number of insertions around the wild-type (WT) sequence. This results in a library, illustrated in Figure 7, containing fragments of variable lengths, spanning from a minimum of 28 (the length of the WT fragment) to a maximum of 57. To overcome the challenge of handling non-aligned sequences effectively, we opted to integrate a convolutional layer into the input of all our architectural models.

In conclusion, our results support the introduction of a biophysical model designed to probabilistically describe the different phases of the experiment—amplification, selection, and sequencing. This model not only provides a robust and interpretable computational framework but also serves as a highly valuable tool for modeling the intricate mutational landscape characteristic of DMS experiments, particularly in the quest for selecting viable AAV2 capsids.

## ACKNOWLEDGMENTS

AP, GU, and JFdCD acknowledge funding from European Unions Horizon 2020 research and innovation programme MSCA-RISE-2016 under grant agreement No.734439 INFERNET. AP, and MDL acknowledge financial support from Future Artificial Intelligence Research (FAIR) and Centro Nazionale di Ricerca in High-Performance Computing, Big Data, and Quantum Computing (ICSC) founded by the European Union-Next Generation EU.

