## Supplementary Information for "Unsupervised modeling of mutational landscapes of adeno-associated viruses viability"

### A. SUPPLEMENTARY INFORMATION

#### A.1. *Datasets*

For their experiments, they first constructed a dataset containing all possible single-site mutations and insertions and then tested it (*experiment-1*). They used the outcome of this first experiment to define a score for each single-site mutation, and this score has been used to generate random mutants in three different ways:

1. With $h_i(a)$ being the score of aminoacid $a$ at site $i$ we can define for each site $i$ the Boltzmann distribution $p_i(a) = \frac{e^{h_i(a)/T}}{Z}$ and sample from it one aminoacid per site. The temperature parameter T controls the "bias" induced by the single-site score.

2. For each site an amino acid is sampled with uniform probability if its score exceeds some chosen threshold.

3. Create variants by randomly performing multiple single-site edits (mutations and insertions) and accept them if the sum of the scores for each edit exceeds some chosen threshold.

In addition to the score-designed variants, a collection of random variants was sampled at distances ranging from 2 to 10 from the wild-type sequence (tested in *experiment-2*).

One of their objectives was to investigate how the composition and size of the training set provided to the deep learning models affect their generative capabilities. For this purpose, they created different training sets based on the sampling methodologies described and the distances of the variants from the wild-type sequence. However, our focus was on exploring how the inclusion of a biophysical model could enhance the quality of inference compared to solely feeding sequences into a neural network and addressing a regression problem. Therefore, we consolidated all the sampled sequences into a single training set for each experiment.

After training their machine learning models, the researchers proceeded to generate new enhanced capsids using the following procedure. Initially, they sampled approximately two billion random sequences within a distance range of 5 to 25 from the wild-type sequence. Subsequently, these random sequences were scored and ranked using the trained model, and the top 1000 sequences at each distance were selected as seed sequences for the generation process. Furthermore, the top 100 sequences were subjected to experimental testing.

Starting from these selected seed sequences, a random set of 250 mutations/insertions was performed. These mutated sequences were then scored and ranked once again using the trained models. The 50 sequences with the highest scores were chosen for the next iteration of mutations/insertions. This iterative process was repeated for a total of 20 rounds, and at each distance, a fixed number of the best sequences were selected. The optimal sequences were selected within a distance range of 5 to 29.

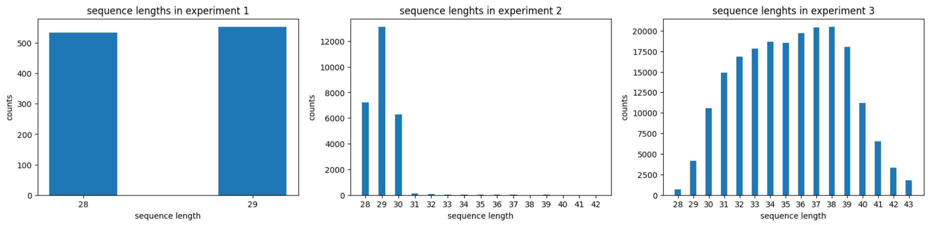


**Figure 8**: Empirical distributions of the distance from the WT sequence of the three data sets *experiment-1,2,3*.

All of these sequences, which were experimentally tested in *experiment-3*, were utilized by us to evaluate the generalization power of the machine learning model. It is worth noting that this seed and generated sequences differ significantly from the ones examined in *experiments 1 and 2*, as the latter were primarily sampled around the wild-type sequence.

#### A.2. *Sequence encoding*

To create diversity in the dataset, sequences have been mutated by substituting the residue present at one site with another one, or by inserting a new residue between two positions. No more than one residue is inserted between two sites. That being said, the mutated sequences have variable lengths ranging from a minimum of 28 (the length of the WT sequence where no insertion has been introduced) to a maximum of 57 (the case in which an insertion has been interposed between each pair of neighbor sites, see Fig. 8 for more details. In order to provide fixed-length input data to the neural network we have chosen the following encoding for sequences. We use a *gap* symbol to pad variants up to the maximum length that can be achieved by the sequences with the peculiar pattern of this dataset (insertion-mutation-insertion-mutation-...-insertion), i.e. 57 as mentioned before. In principle, the side of the sequences to pad would be totally arbitrary. Still, since sequences pass through two regions, one buried inside the capsid which is more conserved, and another one more exposed which is more variable, we decide to put the padding on the side which is more variable since the more conserved sites are more plausible to be aligned.
In this choice we are not consistent with Bryant et al. (2021) because they use an encoding that is equivalent to the following scheme: for even positions put the letter symbol present (WT letter or a mutation), while for odd symbols put a letter symbol if that insertion is performed otherwise put a gap if no insertion is present. We have chosen not to use this latter because if we consider for instance the sequence of letters $A\ -\ B\ C\ D$, it would produce exactly the same sequence given by $A\ B\ C\ -\ D$ by they would have different encodings and therefore they would be different according to the model.
The sequences then are fed to the network as one-hot arrays.

#### A.3. *Architecture of the sequence to energy mapping neural network*

1. Input: the residue sequence of the mutated part, encoded as a one-hot array (see A.2 for details).
2. Convolutional layer: window=7, channels=12, stride=1, activation function=ReLU
3. Batch Normalization (per channel)
4. Pooling layer: type=max, window=2, stride=2

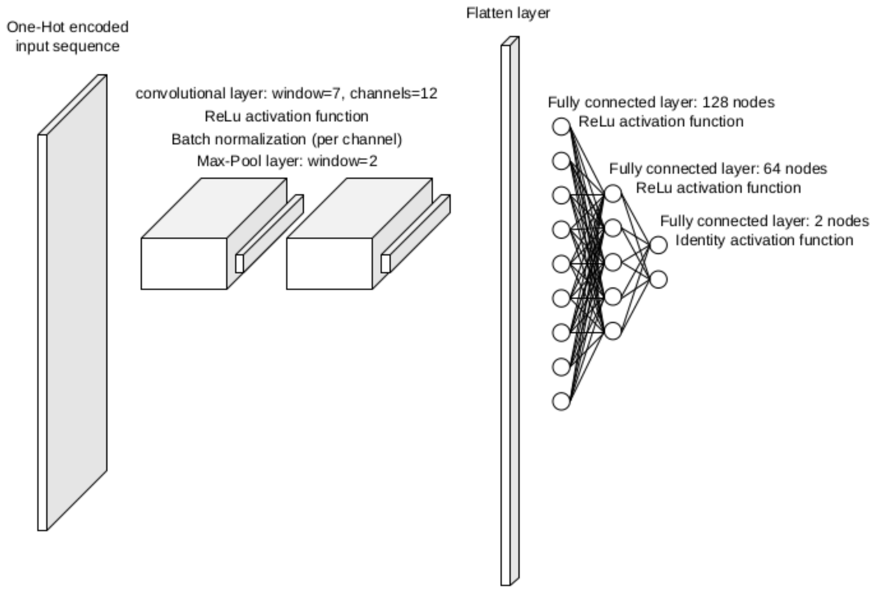


**Figure 9**: Schematic representation of the neural network used in this work. This structure is used both for the standard machine learning approach and for modeling the energy function of the biophysical model. See Sec. A.3 for details on the structure.

5. Convolutional layer: window=7, channels=24, stride=1, activation function=ReLU
6. Batch Normalization (per channel)
7. Pooling layer: type=max, window=2, stride=2
8. Flatten
9. Fully connected: nodes=128, activation function=ReLU.
10. Batch Normalization
11. Fully connected: nodes=64, activation function=ReLU
12. Batch Normalization
13. Fully connected: nodes=2, activation function=identity

Since the energy function of our biophysical models needs to be a number we need to add a further operation to the last layer. To maintain compatibility with the model used in Bryant et al. (2021) we perform the difference between the output of perceptrons in the last layer.

14. difference the output of the last two perceptrons

#### A.4. *Maximum likelihood inference*

As derived in the main-text Eq. (7), the log-likelihood of the model writes:

$$\mathcal{L} = \sum_s R_s \ln\left(\frac{N_s}{\sum_\sigma N_\sigma}\right) + \sum_s R'_s \ln\left(\frac{p_s N_s}{\sum_\sigma p_\sigma N_\sigma}\right) + \text{const.} \tag{A1}$$

which must be maximized over the parameters $\Theta, \mu, N_s$. We first carry out the maximization over $N_s$. Setting the derivative of $\mathcal{L}$ with respect to $N_s$ to zero, we obtain

$$\frac{\partial \mathcal{L}}{\partial N_s} = \frac{R_s + R'_s}{N_s} - \frac{\sum_\sigma R_\sigma}{\sum_\sigma N_\sigma} - \frac{\sum_\sigma p_s R'_\sigma}{\sum_\sigma p_\sigma N_\sigma} = 0 \tag{A2}$$

Solving for $N_s$

$$N_s = \frac{R_s + R'_s}{\frac{\sum_\sigma R_\sigma}{\sum_\sigma N_\sigma} + \frac{\sum_\sigma p_s R'_\sigma}{\sum_\sigma p_\sigma N_\sigma}} \tag{A3}$$

It is convenient to denote the average selectivity of the virus library by

$$\rho = \frac{\sum_s p_s N_s}{\sum_s N_s}, \tag{A4}$$

It then follows that

$$\frac{N_s}{\sum_\sigma N_\sigma} = \frac{\rho(R_s + R'_s)}{\rho \sum_\sigma R_\sigma + \sum_\sigma p_s R'_\sigma} \tag{A5}$$

Note that $\mathcal{L}$ is invariant to scaling of all the $N_s$ by a constant factor. Accordingly, this equation determines the relative abundances $\propto N_s$ only, and not the absolute abundances. The value of $\rho$ must be consistent with its definition (A4), which yields the following self-consistent constraint:

$$\sum_s \frac{p_s(R_s + R'_s)}{\rho \sum_\sigma R_\sigma + \sum_\sigma p_s R'_\sigma} = 1 \tag{A6}$$

Next, for the population after selection and amplification, we find

$$\frac{p_s N_s}{\sum_\sigma p_\sigma N_\sigma} = \frac{p_s(R_s + R'_s)}{\rho \sum_\sigma R_\sigma + \sum_\sigma p_s R'_\sigma} \tag{A7}$$

Substituting (A5) and (A7) into (7), yields

$$\mathcal{L} = \sum_s R_s \ln\left(\frac{\rho(R_s + R'_s)}{\rho \sum_\sigma R_\sigma + \sum_\sigma p_s R'_\sigma}\right) + \sum_s R'_s \ln\left(\frac{p_s(R_s + R'_s)}{\rho \sum_\sigma R_\sigma + \sum_\sigma p_s R'_\sigma}\right) + \text{const.} \tag{A8}$$

Interestingly, we can verify optimizing this expression in $\rho$ results in a stationarity condition equivalent to (A6). Indeed,

$$\begin{aligned} \frac{\partial \mathcal{L}}{\partial \rho} &= \left(\frac{1}{\rho} - \sum_s \frac{R_s + R'_s}{\rho \sum_\sigma R_\sigma + \sum_\sigma p_s R'_\sigma}\right) \sum_s R_s \\ &= \left(1 - \sum_s \frac{\rho(R_s + R'_s)}{\rho \sum_\sigma R_\sigma + \sum_\sigma p_s R'_\sigma}\right) \frac{\sum_s R_s}{\rho} = 0 \end{aligned} \tag{A9}$$

is equivalent to (A6). Therefore, we can treat $\rho$ as a free parameter to be optimized, with condition (A6) being automatically satisfied as soon as the optimum of $\mathcal{L}$ is reached. We therefore, transform our original problem of maximizing $\mathcal{L}(\Theta, \mu, \mathbf{N})$, given by (7), into that of maximizing $\mathcal{L}(\Theta, \mu, \rho)$, given by (A8). This has the advantage of replacing a considerably large number of variables $\mathbf{N} = \{N_s\}$ by the single parameter $\rho$. In practice, we maximize $\mathcal{L}(\Theta, \mu, \rho)$ numerically with the ADAM optimizer Zhuang et al. (2020), and gradients computed by automatic differentiation Innes (2018).
